# Evaluating urban greening scenarios for urban heat mitigation: a spatially-explicit approach

**DOI:** 10.1101/2020.11.09.373779

**Authors:** Martí Bosch, Maxence Locatelli, Perrine Hamel, Rémi Jaligot, Jérôme Chenal, Stéphane Joost

## Abstract

Urban green infrastructure, especially trees, are widely regarded as one of the most effective ways to reducing urban temperatures in extreme heat events, and alleviate its adverse impacts on human health and well-being. Nevertheless, urban planners and decision-makers are still lacking methods and tools to spatially evaluate the cooling effects of urban green spaces and exploit them to assess greening strategies at the urban agglomeration scale. This article introduces a novel spatially-explicit approach to simulate urban greening scenarios by increasing the tree canopy cover in the existing urban fabric, and evaluating their heat mitigation potential. The latter is achieved by applying the InVEST urban cooling model to the synthetic land use/land cover maps generated for the greening scenarios. A case study in the urban agglomeration of Lausanne, Switzerland, illustrates the development of tree canopy scenarios following distinct spatial distribution strategies. The spatial pattern of the tree canopy strongly influences the human exposure to the highest temperatures, and small increases in the abundance of tree canopy cover with the appropriate spatial configuration can have major impacts on human health and well-being. The proposed approach supports urban planning and the design of nature-based solutions to enhance climate resilience.

## Introduction

Urbanization is a global phenomenon that increasingly concentrates the world’s population in urban areas, with the latter expected to grow in both the number of dwellers and spatial extent over the next decades [1–3]. As a major force of landscape change, urbanization is characterized by the conversion of natural to artificial surfaces, which alters the energy and water exchanges as well as the movement of air. Such changes often result in the urban heat island (UHI) effect, a phenomenon by which urban temperatures are warmer than its rural surroundings [4–9]. The negative impacts of UHI have been widely documented and include increased energy and water consumption [10–12], reduced workplace productivity [13, 14] and aggravation of health risks [15–17]. As urban areas grow and global temperatures rise, the UHI effect is expected to become more intense [18, 19], which makes urban heat mitigation a major priority for urban planning and policy-making [20].

Increasing urban green space, especially the urban tree canopy, has been one of the most widely advocated strategies of urban heat mitigation. Nevertheless, the impacts of the urban tree canopy on air temperature show a complex spatial behaviour that remains poorly understood [9, 21, 22]. While many case studies have reported evidence of the cooling effects of urban green areas, the relationship between their size and their cooling capacity is non-linear [23], and little is known about how the overall spatial configuration of urban green spaces affects the heat mitigation at the urban agglomeration scale [24–27]. Therefore, the extent to which cities can use green infrastructure to reduce heat stress remains uncertain, largely because of the lack of fine-grained approaches to evaluate the cooling effects of the spatial pattern of the tree canopy at the urban agglomeration scale.

With the aim of addressing the above shortcomings, the present work introduces a novel spatially-explicit method to evaluate the heat mitigation potential of altering the abundance and spatial configuration of the urban tree canopy cover in realistic settings. The proposed method consists of two major parts. First, synthetic scenarios are generated by increasing the tree canopy cover in candidate locations where the existing urban fabric permits it. Then, the spatial distribution of air temperature of each synthetic scenario is estimated with the InVEST urban cooling model, which simulates urban heat mitigation based on three biophysical processes, namely shade, evapotranspiration and albedo. Finally, the simulated temperature map is coupled with a gridded population census in order to evaluate the human exposure to urban heat in the scenario. By applying such a procedure in the urban agglomeration of Lausanne, Switzerland, this study aims to map the heat mitigation potential that can be achieved starting from the existing urban fabric. With the aim of quantifying the effects of the abundance and spatial configuration of the tree canopy cover on urban heat mitigation, a set of synthetic scenarios are generated by increasing different proportions of tree canopy cover in distinct spatial configurations.

## Materials and Methods

### Study area

Lausanne is the fourth largest Swiss urban agglomeration with 420757 inhabitants as of January 2019 [28]. The agglomeration is located at the Swiss Plateau and on the shore of the Lake Léman, and is characterized by a continental temperate climate with mean annual temperatures of 10.9 °C and mean annual precipitation of 100 mm, with a dominating vegetation of mixed broadleaf forest. The spatial extent of the study has been selected following the recent application of the InVEST urban cooling model to Lausanne by Bosch et al. [29], and covers an area of 112.46 *km*^2^.

In order to evaluate the human exposure to UHI, the population data for the study area has been extracted from the population and households statistics (STATPOP) [30] provided at a 100 m resolution by the Swiss Federal Statistical Office (SFSO) with the Python library swisslandstats-geopy [31].

### Simulation with the InVEST urban cooling model

The spatial distribution of air temperatures is simulated with the InVEST urban cooling model (version 3.8.0) [32], which is based on the heat mitigation provided by shade, evapotranspiration and albedo. The main inputs are a land use/land cover (LULC) raster map, a reference evapotranspiration raster and a biophysical table containing model information of each LULC class of the map. The LULC maps have been obtained by rasterizing the vector geometries of the official cadastral survey of the Canton of Vaud [33] as of August 2019 to a 10 m resolution. Such a dataset distinguishes 25 LULC classes which are relevant ot the urban, rural and wild landscapes encountered in Switzerland. The reference evapotranspiration pixel values are estimated with the Hargreaves equation [34] based on the daily minimum, average and maximum air temperature values of the 1 km gridded inventory of by the Federal Office of Meteorology and Climatology (MeteoSwiss) [35]. The biophysical table used in this study is shown in Table S1. A more thorough description of the model and the data inputs can be found in Bosch et al. [29].

The parameters of the model are set based on its calibration to the same study area in previous work [29]. Finally, the rural reference temperature (*T*_*ref*_) and UHI magnitude (*UHI*_*max*_) values are derived from the air temperature of 11 monitoring stations in the study area (see Figure S1). More precisely, *T*_*ref*_ is set as the 9 p.m. air temperature measurement — the moment of maximal UHI intensity in Switzerland [36] — of the station showing the lowest temperature value, and *UHI*_*max*_ is set as the difference between the 9 p.m. temperature measurement of the station showing the highest temperature value and *T*_*ref*_. With the above definitions, a reference day for the simulations has been selected from the 2018-2019 period as the day showing the maximum *UHI*_*max*_ with *T*_*ref*_ *>* 20. Such a date corresponds to July 27^*th*^ 2018, with *T*_*ref*_ = 20.60°*C* and *UHI*_*max*_ = 7.38°*C*.

### Refining LULC classes based on tree cover and building density

A procedure to redefine the LULC classes from the cadastral survey has been designed to distinguish the LULC classes depending on their proportional cover of both trees and buildings. The reclassification is achieved by combining the 10 m raster LULC map with two 1 m binary raster masks, one for the tree canopy raster and another for the buildings. The 1 m binary tree canopy mask has been derived from the SWISSIMAGE orthomosaic [37], by means of the Python library DetecTree [38], which implements the methods proposed by Yang et al. [39]. The estimated classification accuracy of the tree canopy classification is of 91.75%. On the other hand, the 1 m binary building mask has been obtained by rasterizing the buildings of the vector cadastral survey [33].

The reclassification procedure consists of three steps. Firstly, each 10 m pixel is coupled with the tree canopy and building masks in order to respectively compute its proportion of tree and building cover. Secondly, the set of 10 m pixels of each LULC class are grouped into a user-defined set of bins to form two histograms, one based on their proportion of tree cover and the other analogously for the building cover. Lastly, the two histograms are joined so that each LULC class is further refined into a set of classes. For example, if two bins were used for both the tree and building cover, the “sidewalk” LULC code might be further refined into “sidewalk with low tree/low building cover”, “sidewalk with low tree/high building cover”, “sidewalk with high tree/low building cover” and “sidewalk with high tree/high building cover”.

In the present work, four equally spaced bins (i.e., distinguishing 0-25%, 25-50%, 50-75% and 75-100% intervals) have been used to reclassify each LULC class according to both the tree and building cover. Following the advice given by the directorate of resoures and natural heritage in the Canton of Vaud (DGE-DIRNA), the threshold over which a pixel is considered to have a high tree canopy cover has been set to 75%, which corresponds to placing trees of a spheric crown with a 5 m radius spaced 10 m from one another so that they form a continuous canopy. Therefore, adjacent pixels with a tree canopy cover over 75% can Finally, in order to adapt the biophysical table of the InVEST urban cooling model to the reclassified LULC classes, the shade coefficients are computed as the midpoint of the bin interval of each level of tree cover (i.e., 0.125, 0.375, 0.625 and 0.875), whereas the albedo coefficients have been linearly interpolated based on the level of building cover (see Table S1).

### Generation of urban greening scenarios

Starting from the refined LULC map, a set of urban greening scenarios are generated by altering the LULC classes of certain candidate pixels in a way that corresponds to reasonable transformations that could occcur in urban areas. More precisely, pixels whose base LULC class corresponds to “building”, “road, path”, “sidewalk”, “traffic island”, “other impervious” and “garden” are changed to the LULC code that has the same base class but with the highest tree cover, e.g., pixels of a post-refinement class “sidewalk with low tree/low building cover” are be changed to “sidewalk with high tree/low building cover”. In order to ensure that such an increase of the tree canopy cover is performed only where the existing urban fabric permits it, pixels might only be transformed when two conditions are met. First, the proportion of building cover in the candidate pixels must be under 25%, i.e., there is a 75% of the pixel area which could be occupied by a tree crown. Secondly, pixels of the “road, path” class might only be transformed when they are adjacent to a pixel of a different class, which prevents increasing the tree canopy cover in pixels that are in the middle of a road (e.g., a highway).

After mapping the candidate pixels where the tree canopy cover can be increased, scenarios are generated based on two key attributes: the extent of tree canopy conversion (expressed as a proportion of the total number of candidate pixels), and the selection of pixels to be converted. A set of scenarios is generated by transforming a 12.5, 25, 37.5, 50, 62.5, 75 and 87.5% of the candidate pixels respectively. For each of these canopy areas, three distinct selection approaches are used. The first consists in randomly sampling from the candidate pixels until the desired proportion of changed pixels is matched. In the second and third approaches, the candidate pixels are sampled according to the number of pixels with high tree canopy cover (i.e., greater than 75%) found in their Moore neighborhood (i.e., the 8 adjacent pixels). In the second approach, pixels with higher number of high tree canopy cover neighbors are transformed first, which intends to spatially cluster pixels of high tree canopy cover. The third approach intends to spatially scatter pixels of high tree canopy cover by prioritizing pixels with lower number of high tree canopy neighbors. Given that the three sampling approaches are stochastic, for each scenario configuration, i.e., each pair of proportion of transformed candidate pixels and sampling approach, the corresponding temperature maps will be computed by averaging a number of simulation runs. After observing little variability among the simulation results, the number of runs of a each configuration has been set to 10. Lastly, the set of scenarios is completed with a configuration where a 100% of the candidate pixels are transformed, which is independent of the sampling approach or scenario run since there exists a single deterministic way to transform all the candidate pixels. The final number of scenarios simulated scenarios is 211, i.e., 10 scenario runs for 3 different sampling approaches and 7 proportions of transformed candidate pixels, plus a last scenario where all the pixels are transformed.

For each scenario, the spatial pattern of the tree canopy is quantified by means of a set of spatial metrics from landscape ecology [40, 41], which are computed for the pixels whose post-refinement LULC class has a tree canopy cover over 75% ^1^. Based on other studies that explore the relationship between the spatial of tree canopy and UHIs, four spatial metrics have been chosen to quantify both the composition and oconfiguration of the tree canopy, which are listed in Table 1. The proportion of landscape (PLAND) of pixels with high tree canopy cover serves to quantify the composition aspects, while the configuration is quantified by means of the mean patch size (MPS), edge density (ED) and the mean shape index (MSI) of patches of high tree canopy cover. The four metrics have been computed with the Python library PyLandStats [42].

**Table 1.** Selected landscape metrics. A more thorough description can be found in the documentation of the software FRAGSTATS v4 [41]

| Category | Metric name | Description |
| --- | --- | --- |
| Composition | Percentage of landscape (PLAND) | Percentage of landscape, in terms of area, occupied by pixels with high tree canopy cover |
| Configuration | Mean patch area (AREA_MN) | Average size (in hectares) of the patches formed by pixels with high tree canopy cover |
|  | Mean shape index (SHAPE_MN) | Average shape index of the patches formed by pixels with high tree canopy cover |
|  | Edge density (ED) | Sum of the lengths of all edge segments between pixels with high tree canopy cover an other pixels, per area unit (in m/hectare) |

## Results

### Proportion of transformed pixels by their original LULC class

The relationship between the number of transformed candidate pixels by their original LULC class and the overall proportion of transformed candidate pixels is shown in Figure 1. Changing a 25, 50, 75 and 100% of the candidate pixels corresponds to a total number of pixels changed of 118880, 237760, 356640 and 475520, which account for a total area of 1188.8, 2377.6, 3566.4 and 4755.2 hectares respectively. In the latter case, i.e., increasing the tree canopy in all the possible pixels, 61.50% of the pixels correspond to the “garden” LULC class, followed by “road, path”, “building”, “other impervious”, (18.01, 10.81 and 7.69%, respectively). Finally, the LULC classes of “sidewalk” and “traffic island” constitute only 1.67 and 0.3% of the pixels where the tree canopy can be increased. The differences when considering the sampling approaches separately are small relative to the total number of transformed candidate pixels. The largest differences between sampling approaches can be noted in the number of transformed pixels that originally belong to the “garden” class. When transforming 25, 50 and 75% of the candidate pixels, clustering respectively transforms (on average among the simulation runs) a 0.90, 0.38 and 0.12% more garden pixels than random sampling, and 1.28, 0.76 and 0.43% more garden pixels than the scattering approach (Figure 2).

**Figure 1.**
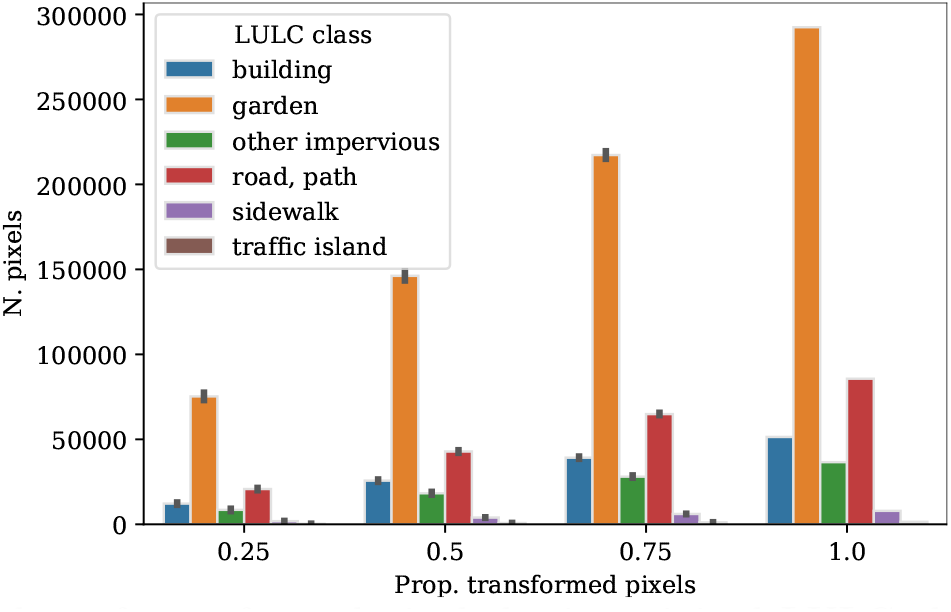
Number of transformed pixels by its original LULC class for an overall proportion of transformed pixels of 25, 50, 75 and 100%. The lines at the top of the bars represent the 95% confidence intervals. The bar heights and the confidence intervals are computed out of all the simulation runs and sampling approaches. See the Jupyter Notebook at section S2.1 for the detailed numbers of the figure.

**Figure 2.**
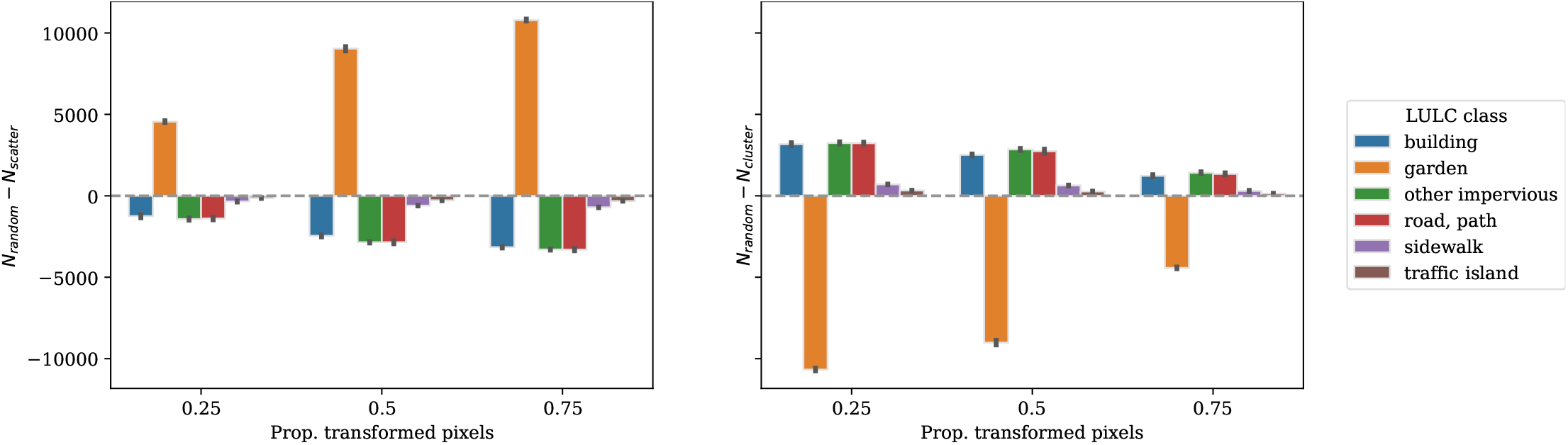
Comparison of between the number of transformed pixels by its original LULC class with the random sampling approach and the scattering (*N*_*random*_ − *N*_*scatter*_, left subplot) and the clustering (*N*_*random*_ − *N*_*cluster*_, right subplot) selection approaches, for an overall proportion of transformed pixels of 25, 50 and 75%. The lines at the top of the bars represent the 95% confidence intervals. The bar heights and the confidence intervals are computed out of all the simulation runs. See the Jupyter Notebook at section S2.1 for the detailed numbers of the figure.

### Simulated LULC, temperature and heat mitigation maps

The LULC, temperature and heat mitigation maps for the scenarios generated by transforming a 25, 50, 75 and 100% of the candidate pixels are shown in Figure 3. When changing 25, 50, 75 and 100% of the candidate pixels, the maximum temperature *T* for the reference date, i.e., 26.05°*C*, is progressively reduced to 25.77, 25.30, 24.82 and 24.49°*C* respectively, while the magnitude of maximum heat mitigation (*T* − *T*_*obs*_) increases from 0.49, 1.17, 1.81 and 2.22°*C* respectively. The largest heat mitigation magnitudes occur in the most urbanized parts, which are located along the main transportation axes. The relationship between the proportion of candidate pixels transformed and the simulated distribution of air temperature can be approximated as a linear relationship with a negative slope (see Figure S2 and Figure S3 for more details about this relationship).

**Figure 3.**
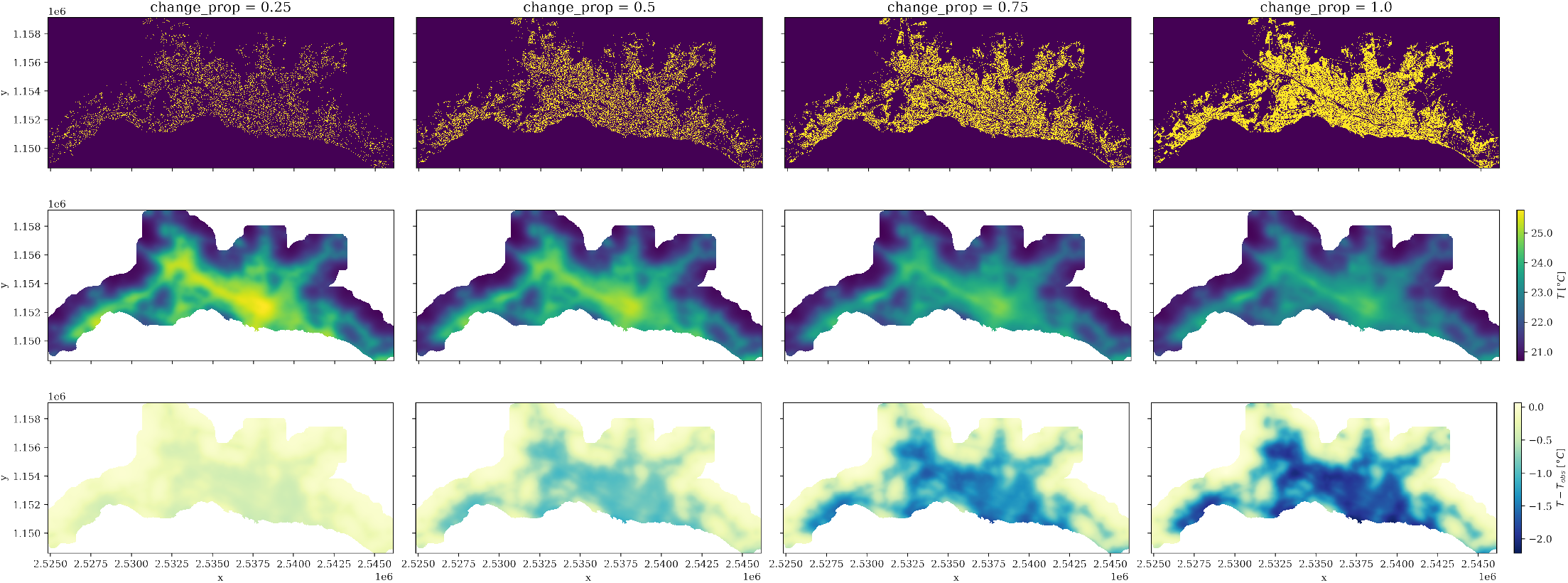
Simulated LULC (top), temperature (middle) and heat mitigation (bottom) maps by transforming a 25, 50, 75 and 100% of the candidate pixels to its corresponding LULC code with high tree canopy cover. The pixel values of each map are aggregated out of all the sampling approaches and simulation runs, i.e., the LULC maps show the mode, whereas the temperature and heat mitigation maps show the average. The axes tick labels display the Swiss CH1903+/LV95 coordinates. See the Jupyter Notebook at section S2.1 for the detailed numbers of the figure.

### Spatial patterns of tree canopy cover

The relationships between the landscape metrics of each scenario run and the corresponding simulated average temperature 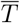 (over all the pixels) are displayed in Figure 4. The proportion of landscape (PLAND) occupied by pixels with high tree canopy cover range from 17.26 to 53.37%. As a composition metric, PLAND is directly related to the proportion of transformed candidate pixels, and the extreme values of the PLAND range correspond to transforming 0 and 100% of the candidate pixels respectively. The relationship between PLAND and the average simulated temperature of each scenario 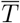 shows a sharp monotonic decrease. However, for the same PLAND values, clustering the transformed pixels to other pixels with high tree canopy cover consistently leads to higher 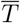 than scattering or randomly sampling — the latter approaches show almost indistinguishable PLAND and 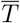 relationship.

**Figure 4.**
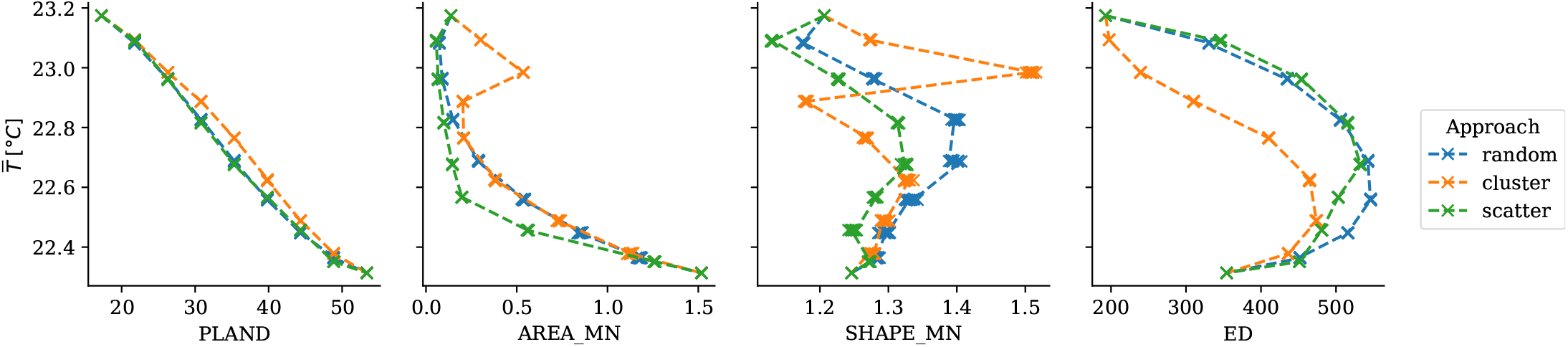
Relationship between landscape metrics and the simulated average temperature 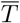 for each scenario run, colored to distinguish the sampling approaches. See the Jupyter Notebook at section S2.2 for the detailed numbers of the figure.

Regarding the configuration metrics, the values of the mean patch area (AREA MN) show that the clustering and random sampling approaches lead to larger patches of high tree canopy cover than the scattering approach. When transforming a 12.5 and 25% of the candidate pixels, clustering them to other pixels of high tree canopy cover increases AREA MN from 0.14 to 0.54 hectares respectively (on average over the simulation runs). For 37.5% of transformed candidate pixels in the clustering approach, AREA MN shows a sudden decline to 0.20 hectares, followed by a monotonic increase that reaches 1.52 hectares when all the candidate pixels are transformed. Such a discernable kink in the computed AREA MN reveals characteristics of the existing urban fabric, and describes the point after which all the candidate pixels that are adjacent to other pixels of high tree canopy have been transformed and hence new pixels have to be allocated as part of new (and smaller) patches. The same kink is even more notable for the mean shape index (SHAPE MN), yet the computed values show a very irregular pattern accross the different scenario configurations, and it is the only metric where differences can be noted among scenario runs with the same configuration. The only consistency is that the scattering approach tends to lower SHAPE MN values than randomly sampling the transformed pixels, which is likely due to the larger abundance of simple single-pixel patches in the former approach. Finally, the clustering approach results in lower edge density (ED) values than in the scattering and random sampling approaches, which show a very similar trend. The observed pattern is consistent with the notion that growing existing patches by clustering the new pixels to them accounts for less total edge length than scattering the same amount of new pixels in a leapfrog manner. In the three approaches, the ED increases monotonically at first until an apex is reached when the proportion of transformed pixels is between 50% and 60%, and then declines monotonically.

The average simulated temperature 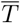 is overall negatively correlated with AREA_MN, which suggests that for the same amount of high tree canopy pixels, large patches provide lower heat mitigation. On the other hand, configurations with the same proportion of high tree canopy pixels show lower 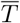 for larger values of ED, which suggests that edge effects between artificial patches and patches of high tree canopy contribute to greater heat mitigation. Nonetheless, as higher proportions of candidate pixels are transformed and the locations of the remaining candidate pixels force the overall ED to decrease, the simulated average temperatures continue to decline. This highlights how the cooling effects of the abundance of tree canopy overshadow those of the spatial configuration, which is consistent with many related research.

### Effects on human exposure

The relationship between human exposure to air temperatures higher than 21, 22, 23, 24, 25 and 26°C and the proportion of pixels transformed to their respective high tree canopy cover class is shown in Figure 5. The number of dwellers exposed to temperatures higher than 21°C does not show a significant decrease (even when converting all the candidate pixels), whereas for temperatures higher than 22°C, it diminishes from 269254 to 268601, 267683, 266518 and 264125 when the proportion of transformed pixels is of 25, 50, 75 and 100% respectively, which represents a relative share of 97.25, 97.02, 96.69 96.27 and 95.41% of the population of the study area. Such a decline progressively becomes more notable as temperatures increase, e.g., the share of the population exposed to temperatures over 24°C declines from an initial 78.4% to 72.39, 59.57, 37.53 and 11.52% when transforming a 25, 50, 75 and 100% of the candidate pixels respectively. Finally, the share of dwellers exposed to temperatures over 25°C, which is initially of 47.91%, is diminished to a 24.98 and 5.74% when transforming a 25 and 50% of the pixels respectively, and becomes 0 after that, whereas the 2508 dwellers originally exposed to temperatures over 26°C do no longer meet such temperatures after transforming a 25% of the candidate pixels.

**Figure 5.**
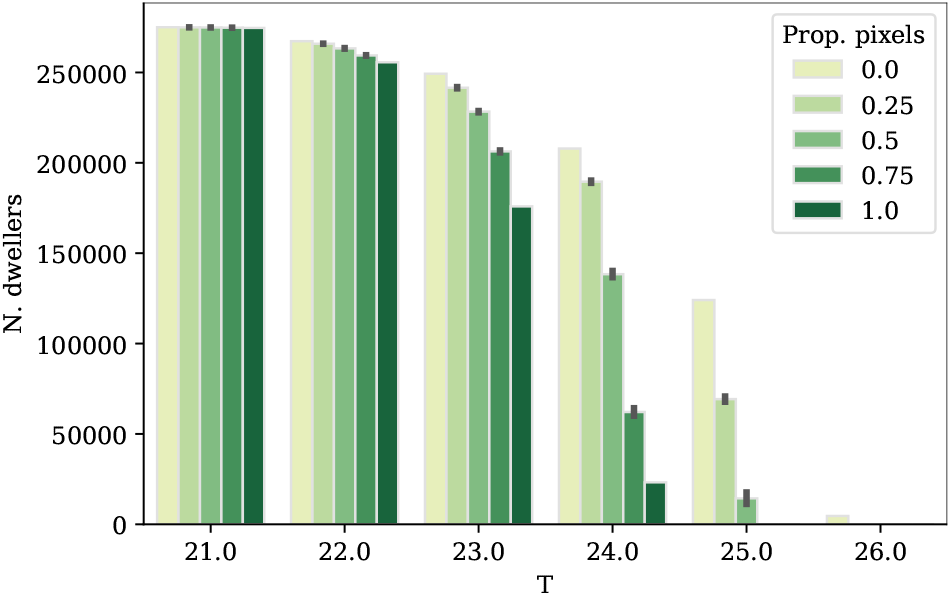
Population exposed to temperatures higher than 21, 22, 23, 24, 25 and 26°C respectively for an overall proportion of transformed pixels of 0, 25, 50, 75 and 100%. The bar heights and the confidence intervals are computed out of all the simulation runs and sampling approaches. See the Juptyer Notebook at section S2.3 for the detailed numbers of the figure.

The way in which the transformed pixels are sampled has significant effects on the human exposure to high temperatures (Figure 6). Overall, scattering the transformed pixels to avoid forming a continuous tree canopy appears as the most effective approach to reduce the human exposure to the highest temperatures, followed by random sampling. When transforming a 25 and 50% of the candidate pixels with the scattering approach, the number of dwellers exposed to temperatures over 25°C decreases from 124073 to 65108 and 4498 respectively. Such a reduction is larger than its random sampling counterpart by 3125 and 8223 dwellers respectively, and larger than its clustering approach counterpart by 9359 and 21388 dwellers respectively (Figure 6).

**Figure 6.**
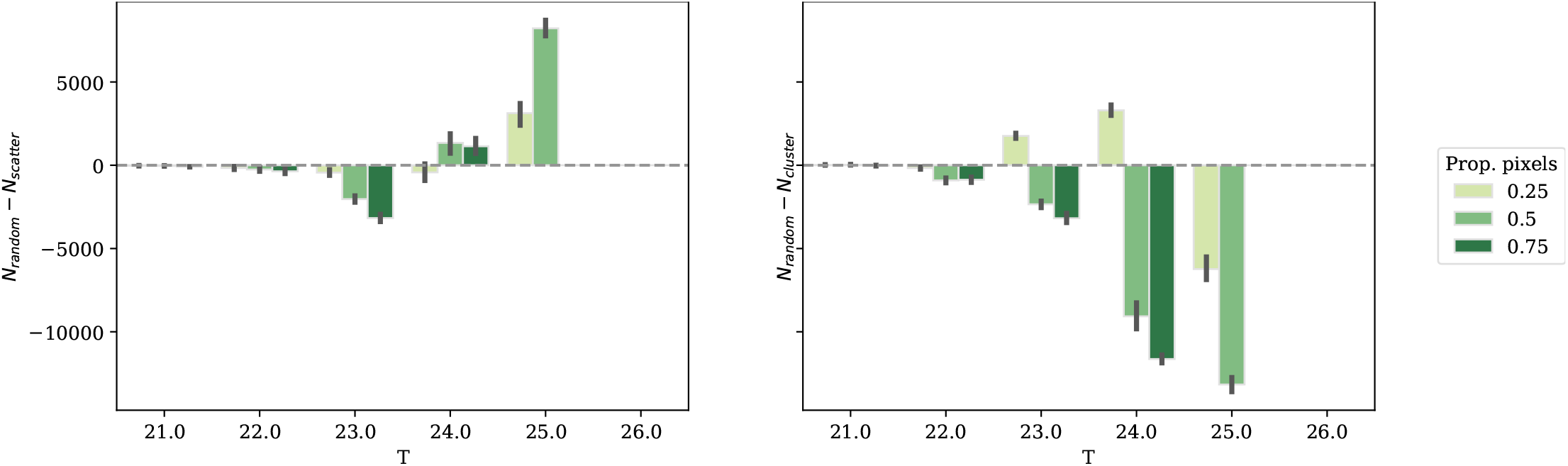
Comparison of between the population exposed to temperatures higher than 21, 22, 23, 24, 25 and 26°C with the random sampling approach and the scattering (*N*_*random*_ − *N*_*scatter*_, left subplot) and the clustering (*N*_*random*_ − *N*_*cluster*_, right subplot) selection approaches, for an overall proportion of transformed pixels of 25, 50 and 75%. The lines at the top of the bars represent the 95% confidence intervals. The bar heights and the confidence intervals are computed out of all the simulation runs. See the Jupyter Notebook at section S2.3 for the detailed numbers of the figure.

## Discussion

### Validity and applicability of the proposed approach

The scenarios simulated in this study map locations where the tree canopy cover in the urban agglomeration of Lausanne can be increased, and suggests that such changes can result in urban nighttime temperatures that are up to 2°C lower. The results indicate that given the same proportion of tree canopy cover, a scattered configuration might lead to more effective urban heat mitigation than a clustered one, which is in line with previous studies in humid climates [43–48]. Nevertheless, the results suggest that effect of the spatial configuration (measured by the metrics AREA_MN, SHAPE_MN and ED) is secondary when compared to the effect of the composition (measured by the PLAND metric). Overall, the effect of the spatial configuration of trees on its urban heat mitigation depends on how it affects the shading and evapotranspiration processes. Such a relationship is known to be strongly mediated by the tree species, background climatic and environmental conditions as well as the spatial scale [45–47, 49–53].

The spatial effects observed in the results are due to the InVEST model equations representing air mixing and the effect of parks. In order to ascertain these effects, the InVEST urban cooling model must be further validated with experiments at the neighborhood scale to ensure that it provides an appropriate city-scale depiction of how the urban heat mitigation mechanisms operate at finer scales. In fact, the InVEST urban cooling model presents limitations regarding the simplified and homogeneous way in which the air is mixed, as well as the cooling effects of large green spaces [29, 32]. As a result, the relationship between the proportion of tree canopy cover and the magnitude of the urban heat mitigation reported in this work is practically linear, and the temperature differences between spatially clustering or scattering the new tree canopy cover are limited. Nonetheless, in complex terrains such as the Lausanne agglomeration, models with uniform weighting of space show considerable deviations from the observed spatial patterns of air temperature [35, 54]. Moreover, the cooling effects of large green spaces have been found to be non-proportional to their area and shape complexity [55–57]. Improving how these non-linear components are represented in the InVEST urban cooling model could enhance not only its validity, but also its value to urban planning by identifying thresholds and regime changes in the cooling efficiency of additional tree planting.

Despite the limitations noted above, a major advantage of the proposed approach is that it can be used to evaluate urban heat mitigation of synthetic scenarios. The simulations presented in this article focus on spatially exploring the effects of an increase of the tree canopy cover, yet there is room for much more experimentation of this kind. On the one hand, the generic sampling approaches explored above can be extended to consider ad-hoc characteristics such as the spatial distribution of the population, and design optimization procedures with specific goals. For instance, the candidate pixels can be selected with the aim of minimizing the exposure of the most vulnerable populations to critical heat thresholds. More broadly, the approach can be used as part of decision support system to explore the trade-offs between ecosystem services provided by trees, perform weighted optimizations and map priority planting locations [58]. On the other hand, in line with recent studies [59–61], the approach could be applied to examine the impact of distinct urbanization scenarios such as densification and urban sprawl on air temperature and human exposure to extreme heat, under current conditions as well as future climate estimates, e.g., by changing the *T*_*ref*_ or *UHI*_*max*_ parameters. Similarly, InVEST urban cooling model might be coupled with models of LULC change such as cellular automata in order to assess not only which scenarios are most desirable in terms of urban heat mitigation, but also which planning strategies might lead to them [62–64].

### Implications for urban planning in Lausanne

The spatiotemporal patterns of LULC change observed during the last 40 years in the Lausanne agglomeration have been characterized by infilling development and a progressive coalescensce of artificial surfaces in its inner ring [65]. Such an infilling trend urges for careful evaluation of the beneficial ecosystem services provided by urban green spaces, which should be balanced against the adverse consequences of urban sprawl [26, 27].

The approach proposed in this study maps locations in the current urban fabric where the tree canopy cover can be increased. While part of this urban greening might occur in impervious surfaces (e.g., in sidewalks, next to roads and in other impervious surfaces), most of the candidate locations currently correspond to urban green space (i.e., the “garden” LULC class). Therefore, the potential heat mitigation suggested by the results study is not attainable in a scenario of severe infill development. Additionally, densification strategies should consider that newly created urban green space might result in less provision of ecosystem services than remnant natural patches [25, 66, 67]. Finally, infilling might exacerbate the unevenness of the accessibility to green areas by depriving dwellers of the most dense parts in city core from their few remaining urban green spaces. Spatial heterogeneity of this kind, which are encountered in many socioeconomic and environmental aspects of contemporary cities, are often hard to represent with aggregate indicators and highlight the importance of spatially explicit models to urban planning and decision making.

The explicit representation of space is also crucial when considering the impacts of urban green space on human exposure to extreme heat. Although the simulated scenarios suggest that the impact of the spatial pattern of tree canopy on the air temperature is practically linear, the implications on human exposure to critical temperatures exhibit important thresholds. For example, by increasing the tree canopy cover of a 25% of the candidate pixels, the number of dwellers exposed to nighttime temperatures over 25°C can be reduced from 124073 to 74466, which respectively represents a 45.08 and 27.06% of the total population in the study area. Furthermore, the results suggest by selecting such pixels to prioritize a spatial scattering of the tree canopy cover, such a population can be reduced by an additional 3125 or 6234 dwellers when respectively compared to random sampling such pixels or clustering them to the existing tree canopy cover. In Switzerland, the excess mortality associated to the heat wave of 2003 occurred over-proportionally to urban and sub-urban residents of its largest urban agglomerations [68]. Furthermore, the association between temperature and mortality in extreme heat events in the largest Swiss urban agglomerations are exponential [69], which indicates that reducing temperatures by even fractions of a degree can have a dramatic impact on death rates.

## Conclusion

The scenarios simulated in this study represent a new way of spatially exploring the heat mitigation potential provided by modifications of the urban fabric, and allow evaluating the cooling effects of both the abundance and spatial configuration of the tree canopy cover. The results map locations where the existing tree canopy cover of the urban agglomeration of Lausanne can be increased, and show an urban cooling potential for urban nighttime temperatures of more than 2°C. Additionally, the simulations suggest that the spatial configuration in which the tree canopy is increased influences its heat mitigation effects. The configuration effects become more significant when considering the impacts on the urban population, and small increases in the tree canopy can result in important reductions in the number of dwellers exposed to the highest temperatures. Overall, the presented approach provides a novel way to explore how the urban tree canopy of can be exploited to reduce heat stress. Future studies can extend the analyses by assessing the provision of other ecosystem services in the various tree canopy strategies presented here.

## Supporting Information

### S1 Data

### S1.1 Biophysical table

The biophysical table for the LULC codes (before the reclassification) is shown in Table S1. The crop and water coefficients are based on Allen et al. [70], while rock, soil and urban coefficients are derived from the results of Grimmond and Oke [71] in the city of Chicago. Given that the evapotranspiration of the vegetation and crops is subject to seasonal changes in temperate zones such as Switzerland [70], the values that correspond to the mid-season estimation (June to August) in [72]. The albedo values are based on the work of Steward et al. [73]. The shade column, which represents the proportion of tree cover of each LULC class, is computed after the reclassification procedure described in section “Refining LULC classes based on tree cover and building density”.

**Table S1.** Biophysical table (before the reclassification). The source comma-separated value (CSV) file used in the computational workflow is available at https://github.com/martibosch/lausanne-heat-islands/blob/master/data/raw/biophysical-table.csv.

| LULC code | Description | Case | $K_c$ | Albedo | Green area |
| --- | --- | --- | --- | --- | --- |
| 0 | building | artificial | 0.4 | 0.1-0.25 | 0 |
| 1 | road, path | artificial | 0.35 | 0.15 | 0 |
| 2 | sidewalk | artificial | 0.35 | 0.15 | 0 |
| 3 | traffic island | artificial | 0.35 | 0.15 | 0 |
| 4 | rail | artificial | 0.35 | 0.15 | 0 |
| 5 | airfield | artificial | 0.4 | 0.2 | 0 |
| 6 | pond | water | 0.45 | 0.15 | 0 |
| 7 | other impervious | artificial | 0.36 | 0.15 | 0 |
| 8 | field, meadow, pasture | vegetation | 0.9 | 0.2 | 1 |
| 9 | vineyards | vegetation | 0.7 | 0.2 | 1 |
| 10 | other intensive farming | vegetation | 1.05 | 0.2 | 1 |
| 11 | garden | artificial | 0.32 | 0.2 | 1 |
| 12 | wetland | water | 0.45 | 0.1 | 1 |
| 13 | other green | vegetation | 0.45 | 0.2 | 1 |
| 14 | backwater | water | 0.65 | 0.05 | 1 |
| 15 | water course | water | 0.65 | 0.05 | 0 |
| 16 | reed | water | 0.45 | 0.1 | 1 |
| 17 | dense forest | vegetation | 1.5 | 0.15 | 1 |
| 18 | densely wooded pasture | vegetation | 1.15 | 0.15 | 1 |
| 19 | open wooded pasture | vegetation | 1.15 | 0.2 | 1 |
| 20 | other wooded | vegetation | 1.15 | 0.15 | 1 |
| 21 | bare rocks | rock and soil | 0.2 | 0.25 | 0 |
| 22 | glacier | water | 0.52 | 0.1 | 0 |
| 23 | sand | rock and soil | 0.3 | 0.25 | 0 |
| 24 | gravel pit | artificial | 0.36 | 0.25 | 0 |
| 25 | other non-vegetated | artificial | 0.36 | 0.15 | 0 |

#### S1.2 Monitoring stations

The locations of the monitoring stations used to get the *T*_*ref*_ and *UHI*_*max*_ parameters of the InVEST urban cooling model are shown in Figure S1. The operators of the stations are: Agrometeo, Federal roads office (ASTRA), Federal office for the environment (BAFU), General directorate for the environment of the Canton of Vaud (DGE), and the Federal Institute of Forest, Snow and Landscape Research (WSL) [74]. The source CSV file with the operator, location and elevation in meters above sea level of the monitoring stations used in the computational workflow is available at https://github.com/martibosch/lausanne-greening-scenarios/blob/master/data/raw/tair-stations/station-locations.csv. The code to produce Figure S1 is available as a Jupyter Notebook (IPYNB) at https://github.com/martibosch/lausanne-greening-scenarios/blob/master/notebooks/stations.ipynb.

**Figure S1.**
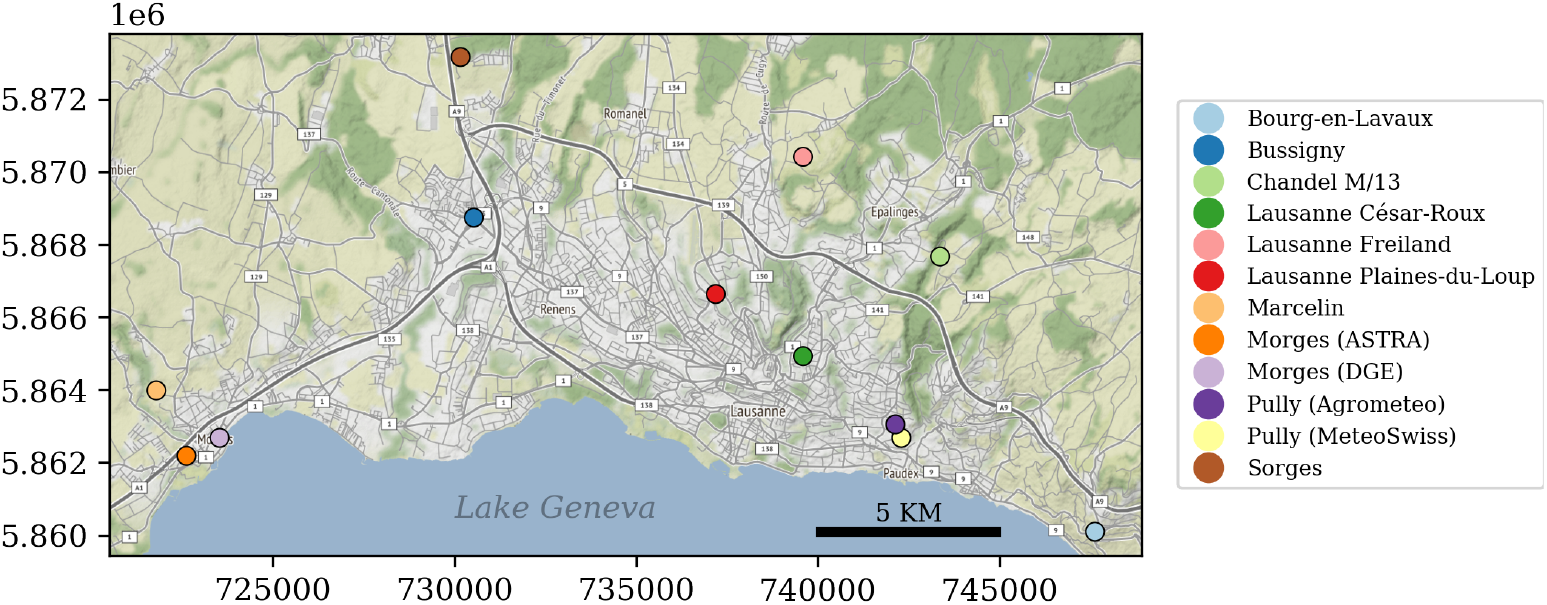
Locations of the monitoring stations used to get the *T*_*ref*_ and *UHI*_*max*_ parameters. The axes tick labels display the Swiss CH1903+/LV95 coordinates. The basemap tile is provided by StamenDesign, under CC BY 3.0, with data from OpenStreetMap, under ODbL.

### S2 Results

#### S2.1 Scenario LULC, temperature and heat mitigation

The code to produce the figures 1, 3, S2 and S3, as well as tables describing the data of the figures, are available as a Jupyter Notebook (IPYNB) at https://github.com/martibosch/lausanne-greening-scenarios/blob/master/notebooks/scenarios.ipynb.

**Figure S2.**
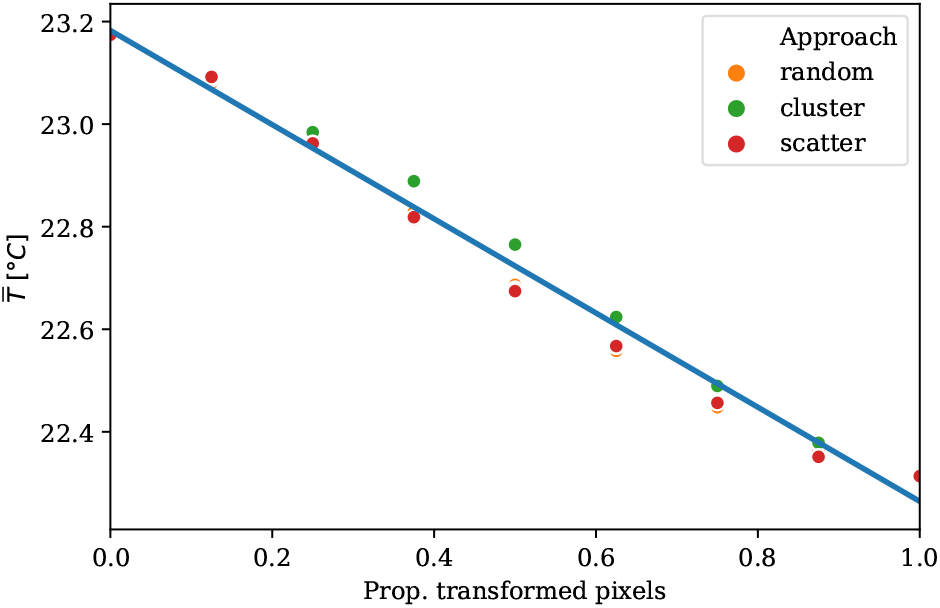
Relationship between the proportion of candidate pixels transformed and the average simulated temperature 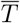 for each scenario sample. The translucent bands around the regression line represent the 95% confidence intervals estimated using a bootstrap.

**Figure S3.**
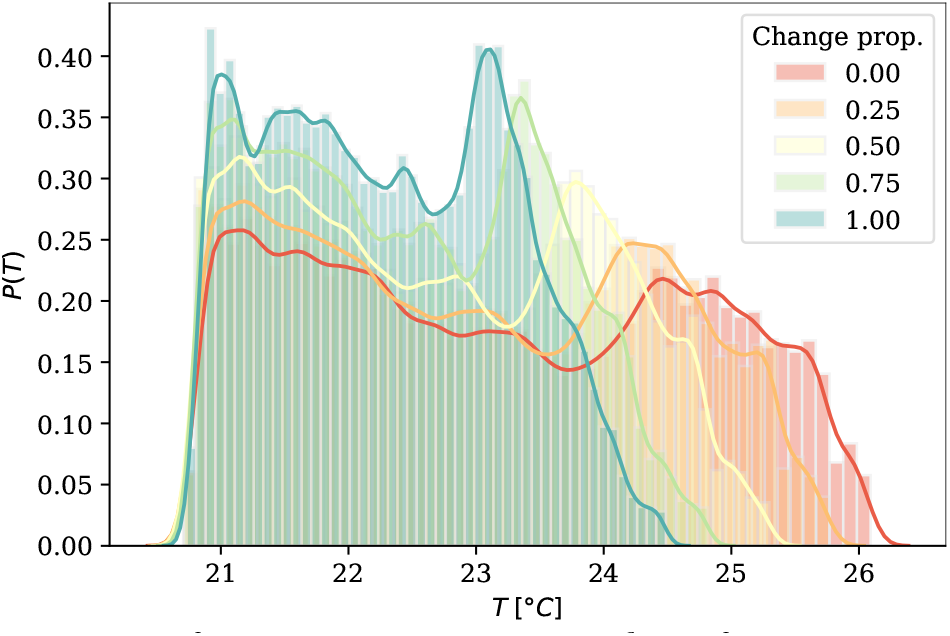
Histogram of raster temperature values for a 25, 50, 75 and 100% of the candidate pixels transformed. The temperature rasters for each histogram are computed by averaging the 10 simulations with the same proportion of candidate pixels transformed.

#### S2.2 Scenario metrics

The code to produce Figure 4 is available as a Jupyter Notebook (IPYNB) at https://github.com/martibosch/lausanne-greening-scenarios/blob/master/notebooks/scenario-metrics.ipynb.

#### S2.3 Scenario human exposure

The code to produce Figure 5 is available as a Jupyter Notebook (IPYNB) at https://github.com/martibosch/lausanne-greening-scenarios/blob/master/notebooks/human-exposure.ipynb.

## Acknowledgments

This research has been supported by the École Polytechnique Fédérale de Lausanne (EPFL).

## Footnotes

1 Following the advice given by the directorate of resoures and natural heritage in the Canton of Vaud (DGE-DIRNA), the threshold over which a pixel is considered to have a high tree canopy cover has been set to 75%, which corresponds to placing trees of a spheric crown with a 5 m radius spaced 10 m from one another so that they form a continuous canopy.

## Notes

https://github.com/martibosch/lausanne-greening-scenarios

### Competing Interest Statement

The authors have declared no competing interest.

## References

1. Karen C Seto, Michail Fragkias, Burak Güneralp, and Michael K Reilly. A meta-analysis of global urban land expansion. PloS one, 6(8):e23777, 2011.

2. Shlomo Angel, Alejandro M Blei, Daniel L Civco, and Jason Parent. Atlas of urban expansion. Lincoln Institute of Land Policy Cambridge, MA, 2012.

3. United Nations. World urbanization prospects: The 2018 revision, 2018.

4. Tim R Oke. City size and the urban heat island. Atmospheric Environment (1967), 7(8):769–779, 1973.

5. Timothy R Oke. The energetic basis of the urban heat island. Quarterly Journal of the Royal Meteorological Society, 108(455):1–24, 1982.

6. A John Arnfield. Two decades of urban climate research: a review of turbulence, exchanges of energy and water, and the urban heat island. International Journal of Climatology: a Journal of the Royal Meteorological Society, 23(1):1–26, 2003.

7. James A Voogt and Tim R Oke. Thermal remote sensing of urban climates. Remote sensing of environment, 86(3):370–384, 2003.

8. Sue UE Grimmond. Urbanization and global environmental change: local effects of urban warming. Geographical Journal, 173(1):83–88, 2007.

9. Patrick E Phelan, Kamil Kaloush, Mark Miner, Jay Golden, Bernadette Phelan, Humberto Silva III, and Robert A Taylor. Urban heat island: mechanisms, implications, and possible remedies. Annual Review of Environment and Resources, 40:285–307, 2015.

10. Hashem Akbari, Melvin Pomerantz, and Haider Taha. Cool surfaces and shade trees to reduce energy use and improve air quality in urban areas. Solar energy, 70(3):295–310, 2001.

11. Jay S Golden, Anthony Brazel, Jennifer Salmond, and David Laws. Energy and water sustainability: The role of urban climate change from metropolitan infrastructure. Journal of Green Building, 1(3):124–138, 2006.

12. Matheos Santamouris, Constantinos Cartalis, Afroditi Synnefa, and Dania Kolokotsa. On the impact of urban heat island and global warming on the power demand and electricity consumption of buildings—a review. Energy and Buildings, 98:119–124, 2015.

13. Tord Kjellstrom, Ingvar Holmer, and Bruno Lemke. Workplace heat stress, health and productivity–an increasing challenge for low and middle-income countries during climate change. Global health action, 2(1):2047, 2009.

14. Kerstin K Zander, Wouter JW Botzen, Elspeth Oppermann, Tord Kjellstrom, and Stephen T Garnett. Heat stress causes substantial labour productivity loss in Australia. Nature Climate Change, 5(7):647–651, 2015.

15. Lauraine G Chestnut, William S Breffle, Joel B Smith, and Laurence S Kalkstein. Analysis of differences in hot-weather-related mortality across 44 US metropolitan areas. Environmental Science & Policy, 1(1):59–70, 1998.

16. R Sari Kovats and Shakoor Hajat. Heat stress and public health: a critical review. Annu. Rev. Public Health, 29:41–55, 2008.

17. Karine Laaidi, Abdelkrim Zeghnoun, Bénédicte Dousset, Philippe Bretin, Stéphanie Vandentorren, Emmanuel Giraudet, and Pascal Beaudeau. The impact of heat islands on mortality in Paris during the August 2003 heat wave. Environmental health perspectives, 120(2):254–259, 2012.

18. Gerald A Meehl and Claudia Tebaldi. More intense, more frequent, and longer lasting heat waves in the 21st century. Science, 305(5686):994–997, 2004.

19. Kangning Huang, Xia Li, Xiaoping Liu, and Karen C Seto. Projecting global urban land expansion and heat island intensification through 2050. Environmental Research Letters, 14(11):114037, 2019.

20. Davide Geneletti, Chiara Cortinovis, Linda Zardo, and Blal Adem Esmail. Planning for ecosystem services in cities. Springer Nature, 2020.

21. Diana E Bowler, Lisette Buyung-Ali, Teri M Knight, and Andrew S Pullin. Urban greening to cool towns and cities: A systematic review of the empirical evidence. Landscape and urban planning, 97(3):147–155, 2010.

22. Carlos Bartesaghi Koc, Paul Osmond, and Alan Peters. Evaluating the cooling effects of green infrastructure: A systematic review of methods, indicators and data sources. Solar Energy, 166:486–508, 2018.

23. L Zardo, D Geneletti, M Pérez-Soba, and M Van Eupen. Estimating the cooling capacity of green infrastructures to support urban planning. Ecosystem services, 26:225–235, 2017.

24. Brenda B Lin and Richard A Fuller. Sharing or sparing? how should we grow the world’s cities? Journal of Applied Ecology, 50(5):1161–1168, 2013.

25. Chi Yung Jim. Sustainable urban greening strategies for compact cities in developing and developed economies. Urban Ecosystems, 16(4):741–761, 2013.

26. Christine Haaland and Cecil Konijnendijk van Den Bosch. Challenges and strategies for urban green-space planning in cities undergoing densification: A review. Urban forestry & urban greening, 14(4):760–771, 2015.

27. Martina Artmann, Luis Inostroza, and Peilei Fan. Urban sprawl, compact urban development and green cities. how much do we know, how much do we agree? Ecological Indicators, 96:3–9, 2019.

28. Swiss Federal Statistical Office. City statistics (urban audit). Data collection. Available from https://www.bfs.admin.ch/bfs/en/home/statistics/ cross-sectional-topics/city-statistics.html. Accessed: 16 April 2019, 2018.

29. Martí Bosch, Maxence Locatelli, Perrine Hamel, Roy P. Remme, Jérôme Chenal, and Stéphane Joost. A spatially-explicit approach to simulate urban heat islands in complex urban landscapes. Geoscientific model development, 2020.

30. Swiss Federal Statistical Office. Statistique de la population et des ménages (statpop), géodonnées 2019. Available from (in French) https://www.bfs.admin.ch/bfs/fr/home/services/geostat/geodonnees-statistique-federale/batiments-logements-menages-personnes/population-menages-depuis-2010.assetdetail.14027479.html. Accessed: 23 September 2020, 2020.

31. Martí Bosch. swisslandstats-geopy: Python tools for the land statistics datasets from the Swiss Federal Statistical Office. Journal of Open Source Software, 4(41):1511, 2019.

32. Richard Sharp, HT Tallis, T Ricketts, AD Guerry, SA Wood, R Chaplin-Kramer, E Nelson, D Ennaanay, S Wolny, N Olwero, et al. Invest 3.8.0 user’s guide. the natural capital project, 2020.

33. Association pour le Système d’information du Territoire Vaudois. Structure des données cadastrales au format shape (md.01-mo-vd). Available from (in French) https://www.asitvd.ch/md/508. Accessed: 3 September 2019, 2018.

34. George H Hargreaves and Zohrab A Samani. Reference crop evapotranspiration from temperature. Applied engineering in agriculture, 1(2):96–99, 1985.

35. Christoph Frei. Interpolation of temperature in a mountainous region using nonlinear profiles and non-euclidean distances. International Journal of Climatology, 34(5):1585–1605, 2014.

36. Burgstall, Annkatrin. Representing the urban heat island effect in future climates. Scientific Report MeteoSwiss, No. 105, 2018.

37. Federal Office of Topography. Swissimage la mosäique d’ortophotos de la Suisse. Available from (in French) https://shop.swisstopo.admin.ch/fr/products/ images/ortho_images/SWISSIMAGE. Accessed: 3 March 2020, 2019.

38. Martí Bosch. Detectree: Tree detection from aerial imagery in Python. Journal of Open Source Software, 5(50):2172, 2020.

39. Lin Yang, Xiaqing Wu, Emil Praun, and Xiaoxu Ma. Tree detection from aerial imagery. In Proceedings of the 17th ACM SIGSPATIAL International Conference on Advances in Geographic Information Systems, pages 131–137. ACM, 2009.

40. R. O’Neill, JR Krummel, RH Gardner, G Sugihara, B Jackson, DL DeAngelis, BT Milne, Monica G Turner, B Zygmunt, SW Christensen, VH Dale, and RL Graham. Indices of landscape pattern. Landscape ecology, 1(3):153–162, 1988.

41. Kevin McGarigal, Sam A Cushman, and Eduard Ene. Fragstats v4: spatial pattern analysis program for categorical and continuous maps. Computer software program produced by the authors at the University of Massachusetts, Amherst. Available from https://www.umass.edu/landeco/research/fragstats/fragstats.html. Accessed: 5 March 2020, 2012.

42. Martí Bosch. PyLandStats: An open-source Pythonic library to compute landscape metrics. PLoS One, 14(12), 2019.

43. Weiqi Zhou, Ganlin Huang, and Mary L Cadenasso. Does spatial configuration matter? understanding the effects of land cover pattern on land surface temperature in urban landscapes. Landscape and urban planning, 102(1):54–63, 2011.

44. Fanhua Kong, Haiwei Yin, Philip James, Lucy R Hutyra, and Hong S He. Effects of spatial pattern of greenspace on urban cooling in a large metropolitan area of eastern China. Landscape and Urban Planning, 128:35–47, 2014.

45. Ronald C Estoque, Yuji Murayama, and Soe W Myint. Effects of landscape composition and pattern on land surface temperature: An urban heat island study in the megacities of Southeast Asia. Science of the Total Environment, 577:349–359, 2017.

46. Weiqi Zhou, Jia Wang, and Mary L Cadenasso. Effects of the spatial configuration of trees on urban heat mitigation: A comparative study. Remote Sensing of Environment, 195:1–12, 2017.

47. Zhaowu Yu, Shaobin Xu, Yuhan Zhang, Gertrud Jørgensen, and Henrik Vejre. Strong contributions of local background climate to the cooling effect of urban green vegetation. Scientific reports, 8(1):1–9, 2018.

48. Mojca Nastran, Milan Kobal, and Klemen Eler. Urban heat islands in relation to green land use in European cities. Urban Forestry & Urban Greening, 37:33–41, 2019.

49. Xiaoma Li, Weiqi Zhou, and Zhiyun Ouyang. Relationship between land surface temperature and spatial pattern of greenspace: What are the effects of spatial resolution? Landscape and Urban Planning, 114:1–8, 2013.

50. Min Jiao, Weiqi Zhou, Zhong Zheng, Jia Wang, and Yuguo Qian. Patch size of trees affects its cooling effectiveness: A perspective from shading and transpiration processes. Agricultural and Forest Meteorology, 247:293–299, 2017.

51. Jingli Yan, Weiqi Zhou, and G Darrel Jenerette. Testing an energy exchange and microclimate cooling hypothesis for the effect of vegetation configuration on urban heat. Agricultural and Forest Meteorology, 279:107666, 2019.

52. Jia Wang, Weiqi Zhou, Min Jiao, Zhong Zheng, Tian Ren, and Qiming Zhang. Significant effects of ecological context on urban trees’ cooling efficiency. ISPRS Journal of Photogrammetry and Remote Sensing, 159:78–89, 2020.

53. Berhanu Keno Terfa, Nengcheng Chen, Xiang Zhang, and Dev Niyogi. Spatial configuration and extent explains the urban heat mitigation potential due to green spaces: Analysis over Addis Ababa, Ethiopia. Remote Sensing, 12(18):2876, 2020.

54. Sylvain Labedens, Jean-Louis Scartezzini, and Dasaraden Mauree. Modeling of mesoscale phenomena using WRF-BEP-BEM-CIM in a complex region. In Journal of Physics: Conference Series, volume 1343, page 012012. IOP Publishing, 2019.

55. Ailian Chen, X Angela Yao, Ranhao Sun, and Liding Chen. Effect of urban green patterns on surface urban cool islands and its seasonal variations. Urban forestry & urban greening, 13(4):646–654, 2014.

56. Tongliga Bao, Xueming Li, Jing Zhang, Yingjia Zhang, and Shenzhen Tian. Assessing the distribution of urban green spaces and its anisotropic cooling distance on urban heat island pattern in Baotou, China. ISPRS International Journal of Geo-Information, 5(2):12, 2016.

57. Hongyu Du, Wenbo Cai, Yanqing Xu, Zhibao Wang, Yuanyuan Wang, and Yongli Cai. Quantifying the cool island effects of urban green spaces using remote sensing data. Urban Forestry & Urban Greening, 27:24–31, 2017.

58. EW Bodnaruk, CN Kroll, Y Yang, S Hirabayashi, DJ Nowak, and TA Endreny. Where to plant urban trees? a spatially explicit methodology to explore ecosystem service tradeoffs. Landscape and Urban Planning, 157:457–467, 2017.

59. Aude Lemonsu, Vincent Viguie, M Daniel, and V Masson. Vulnerability to heat waves: Impact of urban expansion scenarios on urban heat island and heat stress in Paris (France). Urban Climate, 14:586–605, 2015.

60. Long Yang, Dev Niyogi, Mukul Tewari, Daniel Aliaga, Fei Chen, Fuqiang Tian, and Guangheng Ni. Contrasting impacts of urban forms on the future thermal environment: example of Beijing metropolitan area. Environmental Research Letters, 11(3):034018, 2016.

61. Heidelinde Trimmel, Philipp Weihs, Stéphanie Faroux, Herbert Formayer, Paul Hamer, Kristofer Hasel, Johannes Laimighofer, David Leidinger, Valéry Masson, Imran Nadeem, et al. Thermal conditions during heat waves of a mid-European metropolis under consideration of climate change, urban development scenarios and resilience measures for the mid-21st century. Meteorologische Zeitschrift, 2019.

62. Elisabete A Silva, Jack Ahern, and Jack Wileden. Strategies for landscape ecology: An application using cellular automata models. Progress in Planning, 70(4):133–177, 2008.

63. R. White, G. Engelen, and I. Uljee. Modeling Cities and Regions as Complex Systems: From Theory to Planning Applications. MIT Press, 2015.

64. Martí Bosch, Jérôme Chenal, and Stéphane Joost. Addressing urban sprawl from the complexity sciences. Urban Science, 3(2):60, 2019.

65. Martí Bosch, Rémi Jaligot, and Jérôme Chenal. Spatiotemporal patterns of urbanization in three Swiss urban agglomerations: insights from landscape metrics, growth modes and fractal analysis. Landscape Ecology, pages 1–13, 2020.

66. Ranhao Sun and Liding Chen. Effects of green space dynamics on urban heat islands: Mitigation and diversification. Ecosystem Services, 23:38–46, 2017.

67. Jing Wang, Weiqi Zhou, Jia Wang, and Yuguo Qian. From quantity to quality: enhanced understanding of the changes in urban greenspace. Landscape Ecology, 34(5):1145–1160, 2019.

68. Leticia Grize, Anke Huss, Oliver Thommen, Christian Schindler, and Charlotte Braun-Fahrlander. Heat wave 2003 and mortality in Switzerland. Swiss Medical Weekly, 135(13-14):200–205, 2005.

69. Martina S Ragettli, Ana M Vicedo-Cabrera, Christian Schindler, and Martin Röösli. Exploring the association between heat and mortality in Switzerland between 1995 and 2013. Environmental research, 158:703–709, 2017.

70. Richard G Allen, Luis S Pereira, Dirk Raes, Martin Smith, et al. Crop evapotranspiration-guidelines for computing crop water requirements - FAO irrigation and drainage paper 56. Fao, Rome, 300(9):D05109, 1998.

71. Sue Grimmond and Tim Oke. Evapotranspiration rates in urban areas. In Proceedings of the Impacts of Urban Growth on Surface Water and Groundwater Quality HS5, pages 235–243. International Association of Hydrological Sciences Publications, 1999.

72. Mircea-Mărgărit Nistor. Mapping evapotranspiration coefficients in the Paris metropolitan area. GEOREVIEW Scientific Annals of Ştefan cel Mare University of Suceava, Geography Series, 26(1):138–153, 2016.

73. Ian D Stewart and Tim R Oke. Local climate zones for urban temperature studies. Bulletin of the American Meteorological Society, 93(12):1879–1900, 2012.

74. Martine Rebetez, Georg von Arx, Arthur Gessler, Elisabeth Graf Pannatier, John L Innes, Peter Jakob, Markéta Jetel, Marlen Kube, Magdalena Nötzli, Marcus Schaub, et al. Meteorological data series from Swiss long-term forest ecosystem research plots since 1997. Annals of Forest Science, 75(2):41, 2018.

